# An *in vivo* platform to identify pathogenic loci

**DOI:** 10.1101/2023.11.01.565153

**Authors:** Shigehiro Yamada, Tiffany Ou, Sibani Nachadalingam, PreMIER Consortium, Shuo Yang, Aaron N. Johnson

## Abstract

Rare genetic disease discovery efforts typically lead to the identification of new disease genes. PreMIER (<u>Pre</u>cision <u>M</u>edicine Integrated <u>E</u>xperimental <u>R</u>esources) is a collaborative platform designed to facilitate functional evaluation of human genetic variants in model systems, and to date the PreMIER Consortium has evaluated over 50 variants in patients with genetic disorders. To understand if *Drosophila* could be used to identify pathogenic disease loci as part of the PreMIER Consortium, we used tissue-specific gene knockdown in the fly as a proof of principle experiment. Tissue-specific knockdown of seven conserved disease genes caused significant changes in viability, longevity, behavior, motor function, and neuronal survival arguing a set of defined assays can be used to determine if a gene of uncertain significance (GUS) regulates specific physiological processes. This study highlights the utility of a tissue-specific knockdown platform in *Drosophila* to characterize GUS, which may provide the first gene-phenotype correlations for patients with idiopathic genetic disorders

## Introduction

A major challenge in treating undiagnosed diseases is that the underlying genetic and cellular pathologies are often unknown. Recent advances in DNA sequencing technologies have made it possible to identify candidate disease genes and sequence variants in a record number of patients. However, screening long candidate lists of genes of uncertain significance (GUS) and variants of uncertain significance (VUS) to identify a contributing mutation remains a significant challenge. Even when a causative link has been firmly established between a gene and a disease, understanding how the variant affects gene function and overall physiology is difficult to determine with bioinformatics approaches alone (Hartin et al., 2020).

In contrast, established experimental paradigms for characterizing sequence variants have been used in the context of model organism genetics for decades. It is increasingly recognized that diagnostic challenges presented by GUS and VUS uncovered in the clinic can be effectively addressed by model organism research (Link and Bellen, 2020). We established the Precision Medicine Integrated Experimental Resources (PreMIER) platform at the Washington University School of Medicine to apply models systems, including *C. elegans*, *Drosophila*, zebrafish, mice, and human embryonic stem cells, toward characterizing VUS and GUS identified in local patients with undiagnosed genetic diseases.

An overarching goal of the PreMIER Consortium is to use *in vivo* screens to identify GUS. We are using *C. elegans* and *Drosophila* as primary, cost-effective screening tools to maximize the number of genes the consortium can screen. Genetic disease modeling depends on experimental perturbations to gene function, and many disease-associated genes have been conserved between humans, *C. elegans*, and *Drosophila*. In addition, functional genomics tools have been available for use in worms and flies for many years. For example, independent consortiums constructed tens of thousands of publicly available transgenic fly lines that express interfering RNAs (RNAi) against *Drosophila* protein coding genes (Dietzl et al., 2007; Ni et al., 2008), and RNAi can simply be expressed in *C. elegans* media to knock down gene expression (Fraser et al., 2000). One advantage to the fly RNAi system is that transgenic RNAi constructs have been developed under the control of an Upstream Activating Sequence (UAS), which is activated by the Gal4 transcription factor. The fly genome does not encode an endogenous Gal4 protein, but hundreds of fly stocks have been developed that express Gal4 under the control of tissue-specific promoters. By simply crossing flies with a Gal4 transgene to flies carrying a UAS-RNAi transgene targeting a candidate locus, the loss of a potentially disease-causing gene can be tested in a tissue-specific manner in just a single generation. The body systems in flies and humans are also largely analogous, and diseases associated with the circulatory, musculoskeletal, and nervous systems have been successfully modeled in flies (Table 1).

**Table 1.** Genetic disorders recently characterized in *Drosophila*.

| **Locus** | **Symptoms** | **Orthologue** | **Assays** | **Reference** |
| --- | --- | --- | --- | --- |
| CDK19 | intellectual disability,<br>developmental delay | *cdk8* | (1), (2) | (Chung et al., 2020) |
| WDR37 | epilepsy<br>developmental delay | *wdr37* | (2), (3), (5) | (Kanca et al., 2019) |
| TBX2 | cardiovascular development<br>skeletal development | *bifid* | (4) | (Liu et al., 2018) |
| IRF2BPL | neurological dysfunction | *pits* | (4) | (Marcogliese et al., 2018) |
| SCAF4 | neurodevelopment | *CG4266* | (2), (3), (5) | (Friedner et al., 2020) |
| CNOT1 | intellectual disability<br>developmental delay | *Not1* | (5) | (Vissers et al., 2020) |
| JAG2 | muscular dystrophy | *Serrate* | (3) | (Coppens et al., 2021) |
| BCAS3 | developmental delay | *rudhira* | (1), (3), | (Hengel et al., 2021) |
| RNF2 | intellectual disability<br>seizures | *Sce* | (1) | (Luo et al., 2021) |
| EBF3 | hypotonia<br>developmental delay | *kn* | (1) | (Chao et al., 2017) |

(1) lethality and longevity, (2) seizure sensitivity, (3) motor function (climbing assay), (4) neurodegeneration (eye) (5) behavior (copulation assay),

The PreMIER Consortium has effectively used *C. elegans* to identify and characterize GUS, but our experience with flies was comparatively limited. A survey of recent, high-impact disease-modeling studies in *Drosophila* identified six commonly used physiological assays that assess viability, longevity, behavior, and motor function (Table 1). To understand if the six physiological assays could be used as a screening platform to characterize a wide array of potential GUS, we identified seven genes from our patient population in which a clear gene-phenotype relationship has been reported in the literature and that have conserved orthologs in the fly. The clinical symptoms associated with the seven genes included progressive loss of developmental skills, sudden onset seizures, intellectual disabilities, and ataxia. We knocked down each of the seven genes in neurons and in muscle throughout development and into adulthood, and assessed physiological phenotypes in adults. Tissue-specific knockdown of each gene caused significant changes in adult physiology in multiple assays. In many instances, the phenotypes of the experimental flies could be correlated with reported patient symptoms. Although we did not observe heightened seizure sensitivity in any of the genetic backgrounds, our study shows that a set of six physiological assays can be used to show a candidate GUS is required for normal viability, longevity, behavior, or motor function.

## Results

### Seven pathogenic genes from the patient population

Clinicians associated with Washington University School of Medicine have identified DNA sequence variants in hundreds of patients with idiopathic disorders. The first step in the PreMIER pipeline is a bioinformatics analysis to assess the potential pathogenicity of the most likely causative variants. At the time this study initiated, over 20 variants had been analyzed by PreMIER from a human genetics perspective. Seven genes associated with the 20 plus variants fit the essential criteria for this proof-of-principle study: 1) the gene was conserved between human and flies, 2) the associated variant was likely pathogenic after computational analysis, and 3) a phenotype-gene relationship was listed on the Online Inheritance in Man (OMIM) database or in ClinVar. Orthologues of the seven disease-associated loci can be classified as previously unstudied in the fly, unstudied in adult flies, or partially studied in adult flies. **ASNS (OMIM 615574; Asparagine synthetase deficiency)**

Compound transheterozygous missense mutations were identified in asparagine synthetase (ASNS) in 13 year-old female with a history of prematurity, microcephaly, epilepsy, spastic diplegia, developmental delays, and intellectual disability. **Asparagine synthetase deficiency** is characterized by microcephaly, severely delayed psychomotor development, progressive encephalopathy, cortical atrophy, and seizure or hyperekplexic activity. The fly orthologue *AsnS* has not been studied.

### ANO3 (OMIM 615034; Dystonia 24)

A potentially pathogenic *de novo* missense mutation in Anoctamin-3 (ANO3) which encodes a calcium activated chloride channel, was identified in an 18 year-old female with progressive loss of developmental skills. Dystonia is a movement disorder characterized by involuntary muscle movement, and Dystonia 24 is focal dystonia of the neck, laryngeal, and upper limb muscles. The fly orthologue *CG6938* has not been studied.

### EPB41L1 (OMIM 614257; Intellectual developmental disorder, autosomal dominant 11)

A *de novo* single nucleotide deletion in Erythrocyte membrane protein band 4.1, like 1, which encodes a membrane protein that mediates interactions between the erythrocyte cytoskeleton and the plasma membrane, was identified in a 5 year-old female with developmental delay and ataxia. Intellectual developmental disorder, autosomal dominant 11 is characterized by global developmental delay, poor overall growth, sometimes with severe feeding difficulties, facial dysmorphism, and distal skeletal anomalies. The fly orthologue *cora* is unstudied in the adult.

### SLC6A2 (OMIM 604715, Orthostatic intolerance)

A *de novo* missense mutation in Solute carrier family 6, member 2 (SLC6A2), which encodes a norepinephrine transporter responsible for reuptake of norepinephrine into presynaptic nerve terminals, was identified in a patient with neurocognitive disorder, behavioral changes, emotional difficulties and anxiety. Orthostatic intolerance is characterized by adrenergic symptoms that occur when an upright posture is assumed. The fly orthologue *DAT* has been studied in the adult and is required for normal movement and behavior.

### ARHGEF15 (ClinVar, Early infantile epileptic encephalopathy with suppression bursts)

A heterozygous single nucleotide deletion in Rho guanine nucleotide exchange factor 15 (ARHGEF15), which encodes a protein that initiates the exchange of GDP with GTP and promote the association of Rho with its effector molecules, was identified in a 3 year-old with sudden onset seizures. Early infantile epileptic encephalopathy with suppression bursts is characterized by the onset of tonic spasms within the first 3 months of life. The fly orthologue *Exn* is unstudied in the adult.

### SLC9A1 (OMIM 616291; Lichtenstein-Knorr syndrome)

An inherited missense mutation in Solute carrier family 9, member 1 (SLC9A1), which encodes a Na+/H+ antiporter, was identified 3 pediatric siblings with varying degrees of developmental delay and autism. Lichtenstein-Knorr syndrome is characterized by severe progressive sensorineural hearing loss and progressive cerebellar ataxia. The fly orthologue *Nhe2* has been studied in the adult where it plays a role in ommatidia patterning.

### VPS13B (OMIM 216550, Cohen syndrome)

Compound transheterozygous missense mutations were identified in vacuolar protein sorting 13, homolog B (VPS13B), which encodes a protein involved in vacuolar sorting, was identified in 5 year-old boy with Cohen syndrome. Cohen syndrome is characterized by facial dysmorphism, microcephaly, truncal obesity, impaired intellectual development, progressive retinopathy, and intermittent congenital neutropenia. The fly orthologue *Vps13B* has not been studied.

### Six assays to assess pathogenicity

A literature search of rare genetic diseases in which gene-phenotype relationships were established through patient DNA sequencing and subsequent modeling in *Drosophila* identified six physiology assays that were used to show a gene or variant is pathogenic (Table 1). The six assays included 1) lethality assays that determine viability during development, 2) longevity assays to assess lifespan, 3) recovery after mechanically-induced seizures, 4) climbing assays to assess adult motor function in response to a mechanical stimulus, 5) neurodegeneration assays assessing changes in highly patterned units of photoreceptor neurons known as ommatidia, and 6) copulation assays to assess adult behavior (Fig. 1A-F; Table 1). The clinical symptoms associated with the genetic diseases we identified included intellectual disability, developmental delay, epilepsy, and seizures as well as cardiovascular and musculoskeletal disorders (Table 1). The wide range of clinical phenotypes in the reported studies substantially overlapped with the neurological symptoms identified in our set of patients, and included non-neural phenotypes suggesting the six physiological assays could be used to model a range of genetic diseases in future studies.

**Figure 1.**
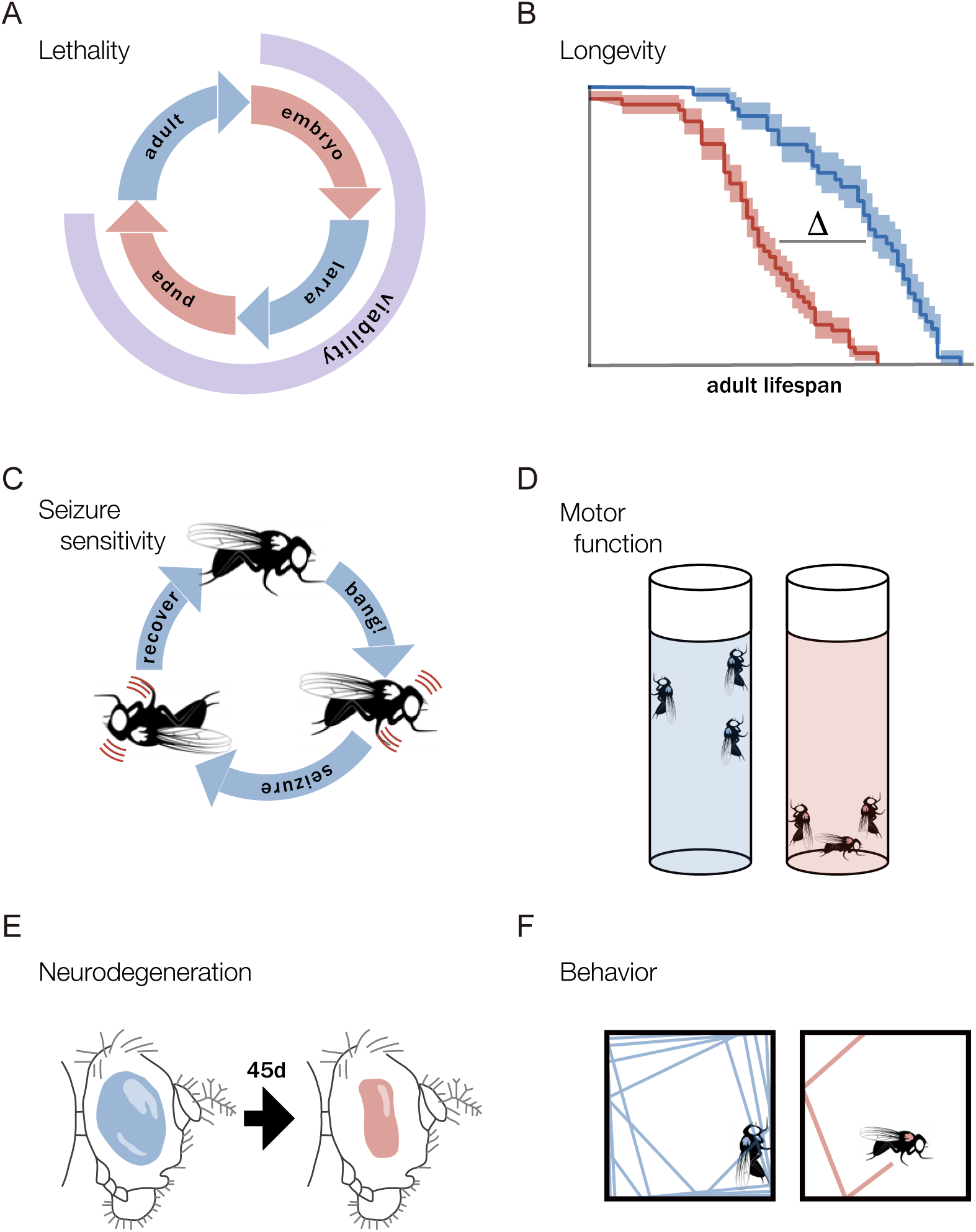
A screening platform to assess tissue-specific gene function *in vivo*. **A.** Lethality assays determine the viability of *elav>RNAi* flies during development. **B.** Longevity assays calculate lifespan. **C.** Seizure sensitivity assays (bang assays) assess recovery in response to vibration-mediated mechanical stimulation. **D.** Climbing assays assess motor function in response to an acute stimulus (tap). **E.** Neurodegeneration assays use ommatidia, which are highly patterned units of photoreceptor neurons, to assess neuron morphology over time. **F.** Open field tests measure voluntary movement in an enclosed arena over time; these assays can reveal behavioral interactions between flies and their environments.

### Viability

We used the pan-neuronal driver *elav.Gal4* to express RNAi constructs against *AsnS, CG6938, cora, DAT, Exn, Nhe2, and Vps13B* (hereafter *elav>RNAi*), and calculated viability for males, females, and both genders combined. *Vps13B^RNAi^*knock down reduced viability in females and combined samples, whereas *elav>DAT^RNAi^*males and *elav>Exn^RNAi^* females showed enhanced viability (Fig. 2A).

**Figure 2.**
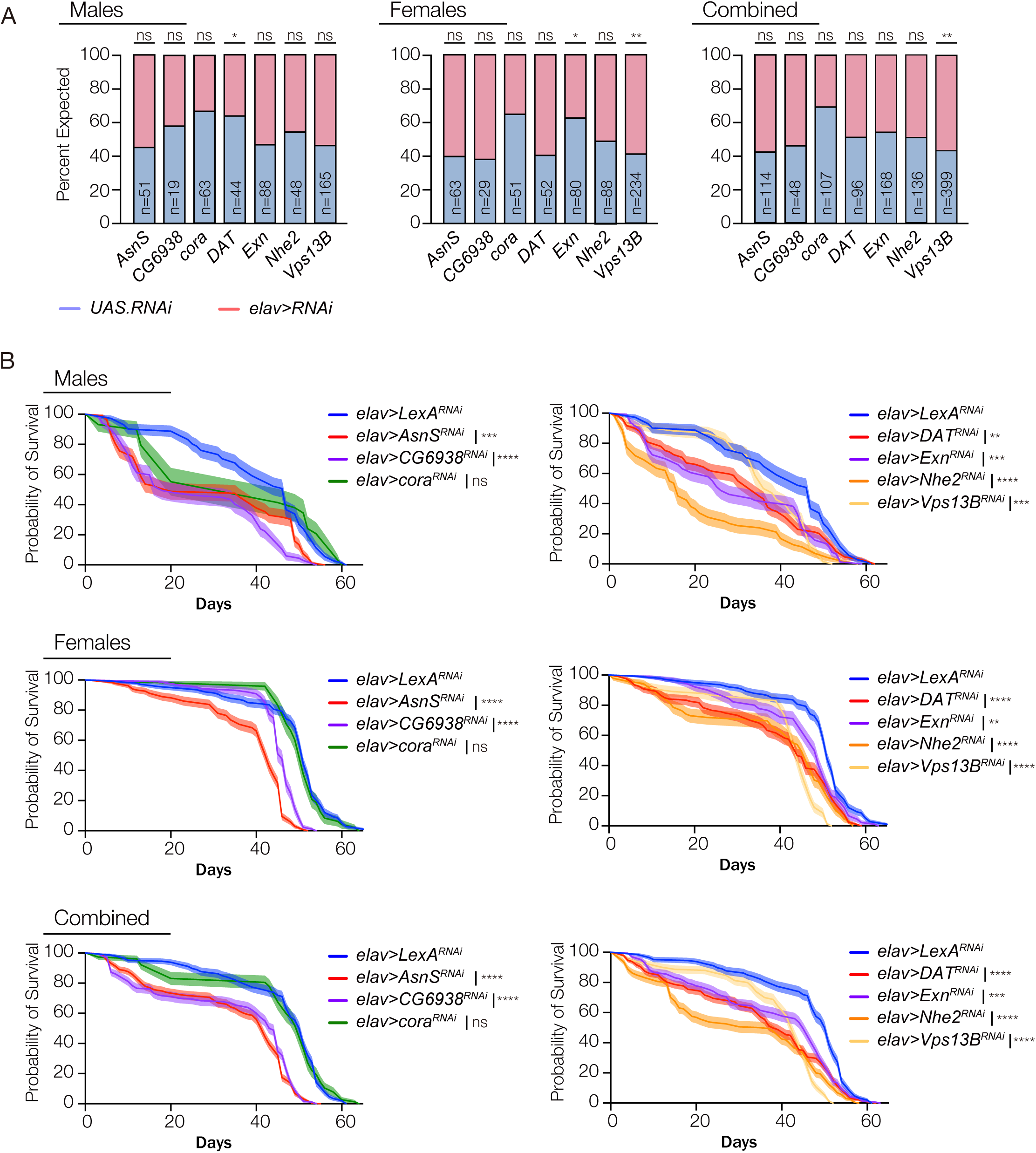
Lethality and longevity assays. **A.** Developmental viability of *elav>RNAi* flies. *elav>DAT^RNAi^* males and *elav>Exn^RNAi^* females showed enhanced viability compared to sibling controls. *elav>Vps13B^RNAi^* females showed a modest yet significant reduction in viability. Note: the expected ratio for *elav>cora^RNAi^*was 1(RNAi):2(control). **B.** Survival curves of *elav>RNAi* flies. With exception of *elav>cora^RNAi^*, all genotypes showed a significant reduction in lifespan compared to *elav>LexA^RNAi^* controls. Males and females did not show differences in longevity. n≥60 flies per cohort. Significance was determined by students unpaired t-test (A) or Kaplan– Meier tests (B). Error bars represent SEM. (*) p< 0.05, (**) p< 0.01, (***) p< 0.001, (****) p < 0.0001, (ns) not significant.

### Longevity

*elav>AsnS^RNAi^, elav>CG6938^RNAi^, elav>DAT^RNAi^, elav>Exn^RNAi^, elav>Nhe2^RNAi^, and elav>Vps13B^RNAi^* males and females showed significantly reduced lifespan compared to *elav>LexA^RNAi^* controls (Fig. 2B). The *Drosophila* genome does encode the *LexA* gene, making *LexA^RNAi^* a suitable control for inducing the RNAi machinery. The lifespan of *elav>cora^RNAi^* flies was comparable to controls (Fig. 2B).

### Seizure recovery

We induced seizures mechanically in *elav>RNAi* flies, and surprisingly found there was no significant change in recovery times among the genotypes or between male and female flies (Fig. 3A).

**Figure 3.**
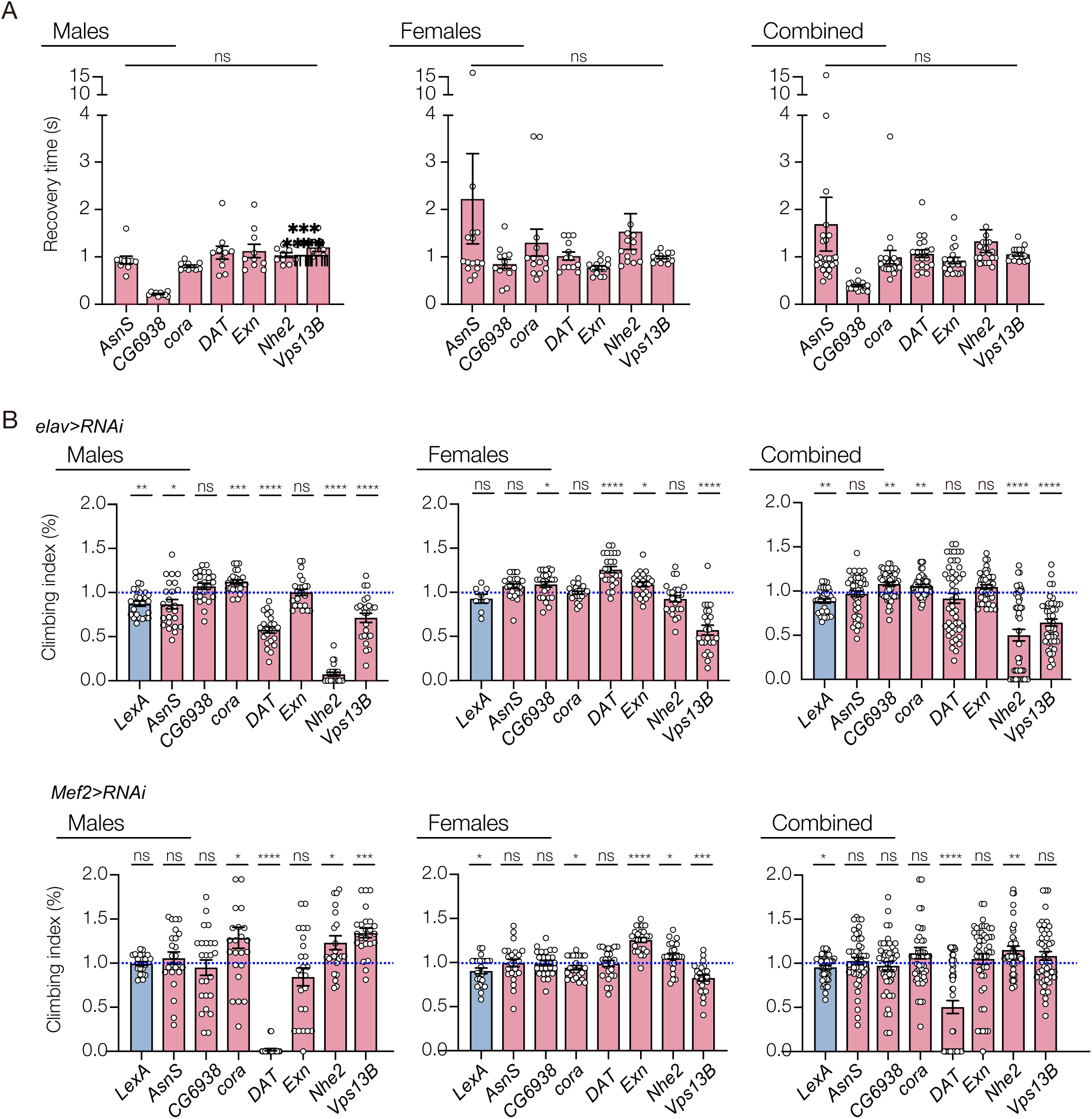
Seizure sensitivity and motor function assays. **A.** Bang assay results of *elav>RNAi* flies. While some *elav>RNAi* individuals showed long recovery times, *elav>RNAi* flies did not show significantly longer recoveries than sibling controls. Each data point represents an individual fly **B.** Climbing assays. *elav>Nhe2^RNAi^* and *elav>Vps13B^RNAi^*flies showed reduced climbing capacity compared to sibling controls. *elav>LexA^RNAi^*flies also showed a modest yet significant reduction in climbing capacity. *elav>AsnS^RNAi^*and *elav>DAT^RNAi^* male flies also showed reduced climbing capacity. Climbing capacity was significantly reduced in *Mef2>DAT^RNAi^* males showing muscle and neuron knockdown of *DAT* can produce similar defects in motor function. Interestingly, *elav>CG6938^RNAi^*, *elav>cora^RNAi^*, and *Mef2>Nhe2^RNAi^* flies showed increased climbing capacity compared to sibling controls. Data points represent the average of individual cohorts, with n=20 flies per cohort. Significance was determined by unpaired student’s t-test. Error bars represent SEM. (*) p< 0.05, (**) p< 0.01, (***) p< 0.001, (****) p < 0.0001, (ns) not significant.

### Motor function

*elav>Nhe2^RNAi^* and *elav>Vps13B^RNAi^* flies showed significantly reduced climbing capacity compared to sibling controls (Fig. 3B). *elav>LexA^RNAi^* flies also showed a slight but significant reduction in climbing capacity, suggesting RNAi alone may have a negative effect on motor function (Fig. 3B). Interestingly, RNAi knockdown differentially affected motor function in male and female flies. For example, climbing capacity was reduced in *elav>AsnS^RNAi^*, *elav>DAT^RNAi^*, and *elav>Nhe2^RNAi^*males flies but was unaffected in female flies (Fig. 3B).

We extended our motor function assays to include muscle-specific knock down using *Mef2.Gal4*. Similar to *elav>DAT^RNAi^* male flies*, Mef2>DAT^RNAi^* males had significantly reduced climbing capacity compared to sibling controls (Fig. 3B). *DAT* encodes a dopamine transporter that mediates dopamine uptake from the synaptic cleft, suggesting DAT is expressed in both motor neurons and myofibers to regulate dopamine levels at the neuromuscular junction. Interestingly muscle-specific knock down of *Nhe2* significantly enhanced climbing capacity, which contrasts the results we observed with neuron-specific knock down of these genes (Fig. 3B). *AsnS* and *Exn* knockdown in either neurons or muscle did not affect motor function.

### Neurodegeneration

To assay neurodegeneration, we used *GMR.Gal4* to express RNAi constructs in photoreceptors neurons after ommatidia cell fates were specified. *GMR>RNAi* ommatidia were collected and imaged in 7- and 45-day old flies. *GMR>CG6938^RNAi^* flies showed a significant increase in the percent of abnormal ommatidia between 7- and 45-days, which included changes in color and morphology indicative of neurodegeneration (Fig. 4A,B). *GMR>cora^RNAi^* flies showed significant ommatidia defects at day 7, that were not further enhanced at day 45 (Fig. 4A,B). *GMR>cora^RNAi^* ommatidia were largely normal at day 1, arguing *cora* knockdown induced neurodegeneration in only six days. Ommatidia morphology was largely unaffected in 7- and 45- day old *GMR>AsnS^RNAi^, GMR>DAT^RNAi^, GMR>Exn^RNAi^*, *GMR>Nhe2^RNAi^,* and *GMR>Vps13B^RNAi^*flies (Fig. 4A,B).

**Figure 4.**
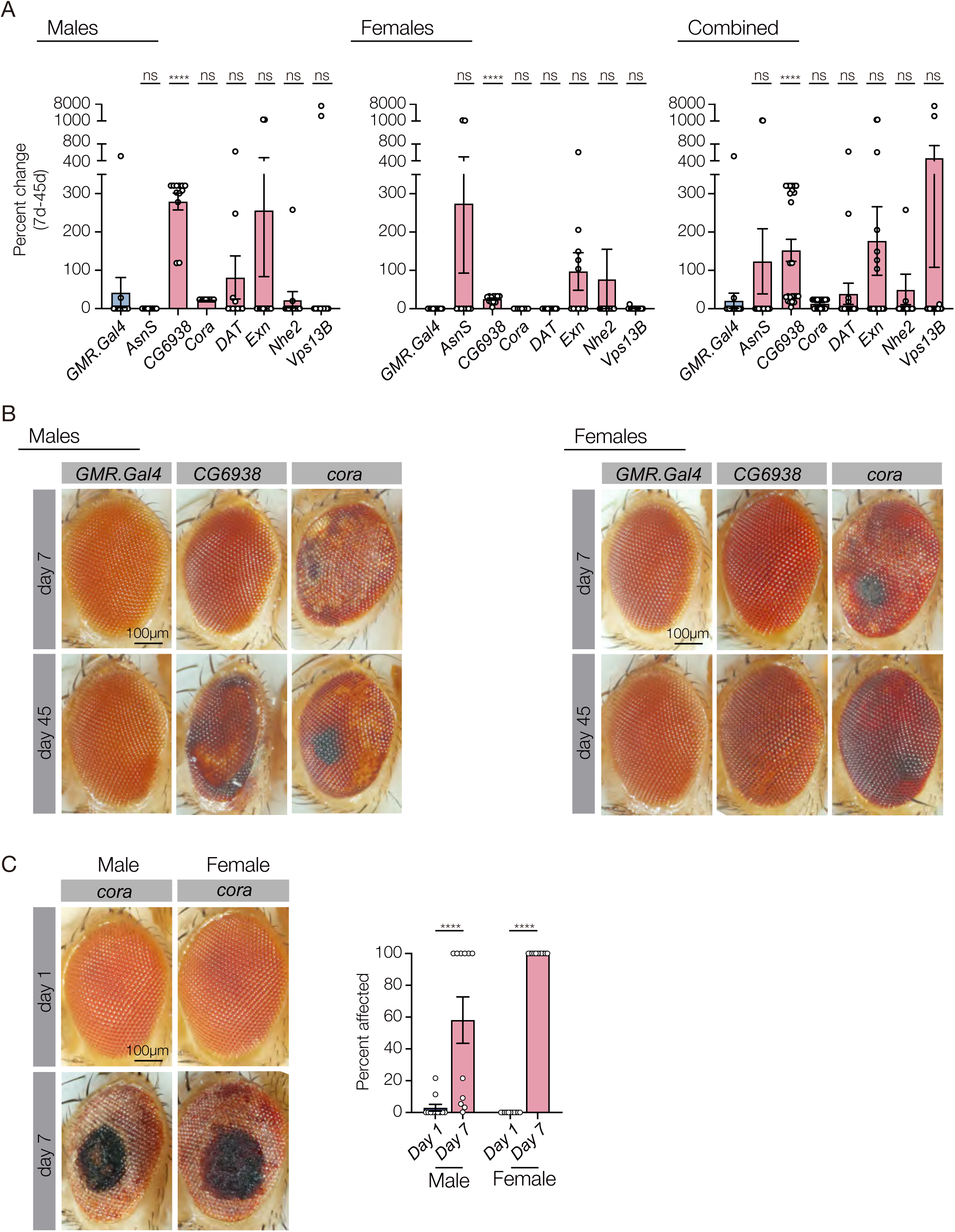
Neurodegeneration assays. **A.** Percent change in abnormal ommatidia between 7 day old and 45 day old *elav>RNAi* flies. *elav>CG6938* flies showed a consistent, significant increase in abnormal ommatidia in both males and females compared to sibling controls. **B.** Micrographs of 7- and 45-day old flies; changes in ommatidia color and morphology where used to score ommatidia as ‘normal’ and ‘abnormal’. Ommatidia in *elav>cora* flies showed morphology defects by day 7, which did not significantly worsen by day 45. **C.** Micrographs of 1- and 7-day old flies; ommatidia in *elav>cora* flies were largely normal at day 1. Since there were no ommatidia defects at day 1, the *elav>cora* phenotype was quantified as percent affected. Error bars represent SEM. (****) p < 0.0001, (ns) not significant.

### Behavior

Copulation assays in *Drosophila* have been used to characterize orthologs of genes associated with developmental delay and epilepsy (Fliedner et al., 2020; Kanca et al., 2019). However open field tests, which measure voluntary movement in an enclosed arena over time, are more commonly used in other species to assess general locomotor activity, exploratory behavior, and emotionality (La-Vu et al., 2020). We used the open field test to assay locomotion (distance traveled), exploration, and anxiety (time immobile). *elav>AsnS^RNAi^* females, *elav>CG6938^RNAi^* males, *elav>cora^RNAi^* flies, *elav>Nhe2^RNAi^* males, and *elav>Vps13B^RNAi^* flies traveled less than sibling controls (Fig. 5A). In addition, *elav>cora^RNAi^* flies, *elav>DAT^RNAi^* flies, and *elav>Vps13B^RNAi^*males explored less than sibling controls (Fig. 5B). *elav>cora^RNAi^*, *elav>DAT^RNAi^*, and *elav>Nhe2^RNAi^*flies were also immobile more often than sibling controls, suggesting these flies had increased anxiety (Fig. 5C). Interestingly, *elav>Exn^RNAi^* males were more active than controls (Fig. 5C). Thus, *elav>RNAi* knockdown of each of the seven disease-associated genes significantly affected one aspect of behavior.

**Figure 5.**
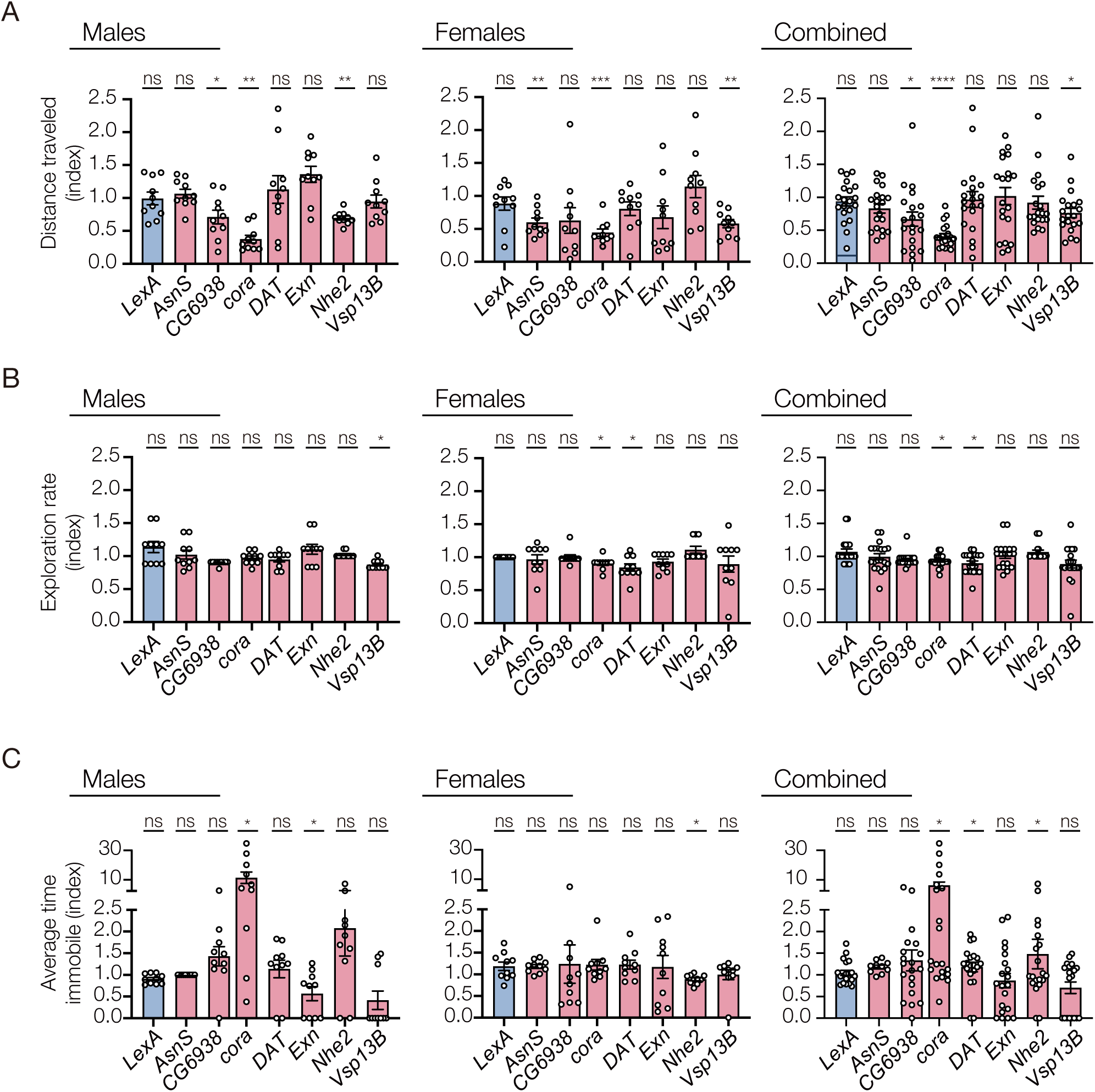
Open field tests. Movement assessments of 3 day old flies for 30 min in behavior arenas. **A.** Distance traveled. *elav>AsnS^RNAi^* (females), *elav>CG6938^RNAi^* (males), *elav>cora^RNAi^* (combined), *elav>Nhe2^RNAi^*(males), and *elav>Vps13B^RNAi^* (combined) flies traveled less than sibling controls. **B.** Exploration rate. *elav>cora^RNAi^* (combined), *elav>DAT^RNAi^* (combined), and *elav>Vps13B^RNAi^*(males) flies explored less than sibling controls. **C.** Time immobile. *elav>cora^RNAi^* (combined), *elav>DAT^RNAi^* (combined), and *elav>Nhe2^RNAi^* (combined) flies were immobile more than sibling controls; *elav>Exn^RNAi^* (males) were immobile less than controls. Data points represent a single individual. Error bars represent SEM. (*) p< 0.05, (**) p< 0.01, (***) p< 0.001, (****) p < 0.0001, (ns) not significant.

## Discussion

Forward genetic approaches in model organisms were the primary means for identifying genes that explained aspects of human biology and disease for decades. Sequencing of the human genome, and more recently the genomes of individual patients, and advances in largescale reverse genetic technologies, such as tissue-specific RNAi, provides a unique opportunity to associate unstudied or understudied genes with clinical phenotypes. The recent explosion in clinical sequencing has unearthed tens of thousands of genes and variants with unknown functions (Frebourg, 2014), which calls for continued reverse genetic approaches in model organisms to validate sequence variants. We have used tissue-specific RNAi against known pathogenic genes to test a battery of physiological assays, and found our approach can largely be applied to any highly-conserved orthologue in *Drosophila* to understand if a gene of uncertain significance (GUS) is required for viability, longevity, motor function, neuron survival, or normal behavior (Table 2).

**Table 2:** Tissue-specific knock down of known pathogenic genes.

| locus<br>human/fly | viability/<br>longevity | seizure<br>recov-<br>ery | motor function |  | neuro-<br>degenera-<br>tion | behavior |  |  |
| --- | --- | --- | --- | --- | --- | --- | --- | --- |
|  |  |  | CNS | muscle |  | locomotion | exploration | anxiety |
| *ASNS/AsnS* | ns/**** | ns | ns | ns | ns | ns | ns | ns |
| *ANO3/CG6938* | ns/**** | ns | ** | ns | **** | * | ns | ns |
| *EPB41L1/cora* | ns/ns | ns | ** | ns | **** | **** | * | * |
| *SLC6A2/DAT* | ns/**** | ns | ns | **** | ns | ns | * | * |
| *ARHGEF15/Exn* | ns/*** | ns | ns | ns | ns | ns | ns | ns |
| *SLC9A1/Nhe2* | ns/**** | ns | **** | ** | ns | ns | ns | * |
| *VPS13B/Vps13B* | **/**** | ns | **** | ns | ns | * | ns | ns |

Results for combined gender analysis shown

### ASNS

The patient from this study with compound transheterozygous variants in *ASNS* had overlapping symptoms with Asparagine synthetase deficiency including microcephaly, seizures, developmental delays, and intellectual disability. *AsnS* knock down in either the CNS or muscle produced relatively mild phenotypes, and only significantly affected longevity (Table 2, Fig. 2B). The human and fly genes have high homology, but paralogous asparagine synthetases in the fly genome may compensate for knock down of *AsnS*. Alternatively, the RNAi construct not produce a strong loss of RNA or protein function.

### ANO3

Symptoms in the patient from this study with a *de novo* missense variant in ANO3 included progressive loss of developmental skills, which overlapped poorly with features of Dystonia 24 (dystonia of the neck, laryngeal, and upper limb muscles). *CG6938* knock down affected longevity, motor function, neuron survival, and voluntary locomotion. Dystonia 24 is thought to arise from abnormal neuron excitability (Charlesworth et al., 2012), but our studies suggest that *ANO3* may also be required for neuron survival.

### EPB41L1

The patient from this study with a *de novo* single nucleotide deletion in *EPB41L1* had overlapping symptoms with *EPB41L1*-associated intellectual developmental disorder, autosomal dominant 11, that include developmental delay and ataxia. *cora* knock down caused defects in motor function after acute stimulation (climbing assay) and reduced voluntary movement in the open field test. Cora is a component of invertebrate septate junctions, which are analogous to vertebrate tight junctions (Fehon et al., 1994). Tight junctions play an essential role in maintaining the blood brain barrier, and in preventing neurodegeneration (Knox et al., 2022). Although EPB41L1 is not known to be a tight junction protein, Cora and EPB41L1 may play analogous roles in promoting cell-cell adhesion and maintaining neuron survival.

### SLC6A2

The patient from this study with a *de novo* missense mutation in *SLC6A2* presented with neurocognitive disorder, behavioral changes, emotional difficulties and anxiety. These symptoms appear to be distinct from *SLC6A2*-associated orthostatic intolerance that mainly describes neurocirculatory symptoms after changes in posture. *SLC6A2* encodes a norepinephrine transporter responsible for reuptake of norepinephrine into presynaptic nerve terminals, whereas the fly orthologue *DAT* encodes a transporter that mediates uptake of dopamine, the norepinephrine precursor, from the synaptic cleft. *DAT* was the only gene that, when knocked down in either the CNS or muscle, caused reduced motor function. This result highlights the fact that tissue-specific knock down tightly correlates with the cellular functions of the targeted proteins, arguing our approach is both specific and highly sensitive.

### ARHGEF15

Symptoms in the patient from this study with a heterozygous single nucleotide deletion in *ARHGEF15* had sudden onset seizures, which largely overlaps with the onset of tonic spasms reported fpr *ARHGEF15-*associated early infantile epileptic encephalopathy with suppression bursts. *Exn* knock down in either the CNS or muscle produced relatively mild phenotypes, and only significantly affected longevity (Table 2, Fig. 2B). *Exn* and *ARHGEF15* are only modestly conserved, and up to 18 *Exn* paralogs have been identified in the fly genome. Our *Exn* results argue mechanical induction of seizures may not be the optimal method to identify seizure-related phenotypes or that *Drosophila* should only be used to model highly orthologous genes.

### SLC9A1

The patients from this study with an inherited missense mutation in *SLC9A1* were diagnosed with developmental delay and autism, which appears to only partially overlap with *SLC9A1*-associated Lichtenstein-Knorr syndrome that is characterized by severe progressive sensorineural hearing loss and progressive cerebellar ataxia. Knock down of *Nhe2* affected longevity and motor function (Table 2, Fig. 2B). Interestingly, the patient from this study showed signs of autism and *Nhe2* knock down flies showed autism-like phenotypes (increased anxiety; Table 2, Fig. 5C). It is possible that deafness and ataxia in Lichtenstein-Knorr syndrome patients may have masked or overshadowed symptoms of autism.

### VPS13B

Compound transheterozygous missense mutations were identified in *VPS13B* in a patient from this study with Cohen syndrome, which is characterized by facial dysmorphism, microcephaly, truncal obesity, impaired intellectual development, progressive retinopathy, and intermittent congenital neutropenia. *Vps13B* knock down affected longevity, motor function, and voluntary movement (Table 2, Figs. 2B, 5A).

## Conclusions

In summary, our *Drosophila* platform can be used to understand if a GUS identified by sequencing patient DNA is required for viability, longevity, motor function, neuronal survival, or normal behavior, provided the orthologues are highly-conserved between flies and humans. Knock down of pathogenic loci in flies often produced phenotypes that overlap with patient symptoms, but phenotypic variability within the patient population substantiates the fact that not all clinical phenotypes need to be observed in flies to substantiate the hypothesis that a given locus is pathogenic. We find it incredibly interesting however that modeling known disease-causing genes with tissue-specific RNAi in can enhance our understanding of gene-phenotype relationships as evidenced by our studies of genes associated with Dystonia 24, orthostatic intolerance, and Lichtenstein-Knorr syndrome. We are also encouraged that once GUS’s are identified in future studies that the genetic tools in *Drosophila* can be applied toward gaining deep biological and mechanistic insights into genetic diseases.

## Materials and methods

### Drosophila genetics

Flies were maintained on standard cornmeal fly food and cultured at 25°C under a normal light/dark cycle. The *Drosophila* stocks were obtained from the Bloomington Stock Center and are described in Flybase (http://flybase.org/) unless otherwise specified. The RNAi lines were *UAS-AsnS-RNAi* (stock 35739), *UAS-CG6938-RNAi* (stock 65941), *UAS-cora-RNAi* (stock 28933), *UAS-DAT-RNAi* (stock 31256), *UAS-Exn-RNAi* (stock 33373), *UAS-Nhe2-RNAi* (stock 51491), *UAS-Vps13B-RNAi* (stock 52915), and *UAS-LexA-RNAi* (stock 67945). Gal4 lines used were *elav-Gal4,Sb/TM6b* (Dr. Helen McNeill, WUSM), *GMR-Gal4* (stock 49924), and *Mef2-Gal4* (stock 50742).

### Lethality assays

30 *elav-Gal4,Sb/TM6b* virgins were crossed to 15 UAS-RNAi males in a bottle. Adults are passed forward every three days. Once the adults began emerging, the number of *Sb(+)* and *Sb(-)* adults were scored daily.

### Longevity assays

1 day old adult flies were collected and transferred to fresh food daily. For longevity analysis, the number of dead flies was recorded daily. Kaplan-Meier survival curves were generated, and statistical analysis was performed using log-rank analysis (Prism9, GraphPad Software).

### Bang assays

To assess seizure sensitivity, a single 3-4 day old adult is transferred to an empty vial, allowed to equilibrate for 1min, and the vortexed for 10s as described ??. The elapsed time for the fly to regain a standing position was evaluated with a timer and recorded as recovery time.

### Climbing assays

To assess motor function, climbing assays were performed as described (Moore et al., 2018). 15-20 flies were placed into empty vials (9.5 cm high,1.7 cm in diameter) with flat bottoms, the flies were forced to the bottom of a vial by firmly tapping the vial against the bench surface. Flies were imaged eight seconds after the tap, and the number of flies that climbed above a 5-cm mark were recorded as positive.

### Neurodegeneration assays

1d old adult flies were collected and transferred to fresh food daily. Flies were collected at 7d and 45d; adults heads were whole mounted and imaged on a AxioZoom V16 (Carl Zeiss). ImageJ software (National Institutes of Health) was used to calculate the percent of ommatidia with normal and abnormal morphology or color. Percent change in abnormal ommatidia from 7d to 45d was calculated to determine degeneration.

### Open field test

3 day old flies were anesthetized and loaded into one cm^2^ arenas, with a thickness of 1.6mm, as described (Mohammad et al., 2016). After equilibrating for 30 min, the flies were video recorded for 30 min. ToxTrac software (Rodriguez et al., 2018) was used to evaluate movement parameters.

### Bioinformatic and statistical analysis

Comparisons of two samples were made using Student’s t-test, and Welch’s correction was applied to samples with unequal standard deviation. Chi-square tests were used to evaluate lethality assays. Survival curves were compared using the Kaplan–Meier test. P values of less than 0.05 were considered statistically significant. All statistical analyses were performed with GraphPad Prism 9 software. The sample sizes and number of replicates are indicated in the figure legends. Data collection and data analysis were routinely performed by different authors to prevent potential bias. All individuals were included in data analysis.

### Ethics statement

Researchers had no access to identifiable patient data. The work included in this manuscript is considered non-human subjects research.

## Acknowledgments

This work was funded by grants from the National Institutes of Health (1R01HD110556) to A.N.J and the PreMIER Consortium.

## Author Contributions

Conceptualization, S.Y., S.H.Y, and A.N.J; Methodology and Validation, S.Y., S.H.Y., and A.N.J.; Formal Analysis, S.Y., S.H.Y, T.O., S.N. and A.N.J.; Investigation, S.Y., S.H.Y., T.O., and S.N.; Writing Original Draft, S.Y. and A.N.J.; Review & Editing, S.Y. and A.N.J.; Supervision, S.Y., and A.N.J.; Funding Acquisition, A.N.J.

## Data availability

The data that support the findings of this study are available from the corresponding authors upon request.

## References

Chao, H. T., Davids, M., Burke, E., Pappas, J. G., Rosenfeld, J. A., McCarty, A. J., Davis, T., Wolfe, L., Toro, C., Tifft, C. et al. (2017). A Syndromic Neurodevelopmental Disorder Caused by De Novo Variants in EBF3. Am J Hum Genet 100, 128–137.

Charlesworth, G., Plagnol, V., Holmström, K. M., Bras, J., Sheerin, U. M., Preza, E., Rubio-Agusti, I., Ryten, M., Schneider, S. A., Stamelou, M. et al. (2012). Mutations in ANO3 cause dominant craniocervical dystonia: ion channel implicated in pathogenesis. Am J Hum Genet 91, 1041–50.

Chung, H. L., Mao, X., Wang, H., Park, Y. J., Marcogliese, P. C., Rosenfeld, J. A., Burrage, L. C., Liu, P., Murdock, D. R., Yamamoto, S. et al. (2020). De Novo Variants in CDK19 Are Associated with a Syndrome Involving Intellectual Disability and Epileptic Encephalopathy. Am J Hum Genet 106, 717–725.

Coppens, S., Barnard, A. M., Puusepp, S., Pajusalu, S., Õunap, K., Vargas-Franco, D., Bruels, C. C., Donkervoort, S., Pais, L., Chao, K. R. et al. (2021). A form of muscular dystrophy associated with pathogenic variants in JAG2. Am J Hum Genet 108, 840–856.

Dietzl, G., Chen, D., Schnorrer, F., Su, K. C., Barinova, Y., Fellner, M., Gasser, B., Kinsey, K., Oppel, S., Scheiblauer, S. et al. (2007). A genome-wide transgenic RNAi library for conditional gene inactivation in Drosophila. Nature 448, 151–6.

Fehon, R. G., Dawson, I. A. and Artavanis-Tsakonas, S. (1994). A Drosophila homologue of membrane-skeleton protein 4.1 is associated with septate junctions and is encoded by the coracle gene. Development 120, 545–557.

Fliedner, A., Kirchner, P., Wiesener, A., van de Beek, I., Waisfisz, Q., van Haelst, M., Scott, D. A., Lalani, S. R., Rosenfeld, J. A., Azamian, M. S. et al. (2020). Variants in SCAF4 Cause a Neurodevelopmental Disorder and Are Associated with Impaired mRNA Processing. Am J Hum Genet 107, 544–554.

Fraser, A. G., Kamath, R. S., Zipperlen, P., Martinez-Campos, M., Sohrmann, M. and Ahringer, J. (2000). Functional genomic analysis of C. elegans chromosome I by systematic RNA interference. Nature 408, 325–330.

Frebourg, T. (2014). The challenge for the next generation of medical geneticists. Hum Mutat 35, 909–11.

Hartin, S. N., Means, J. C., Alaimo, J. T. and Younger, S. T. (2020). Expediting rare disease diagnosis: a call to bridge the gap between clinical and functional genomics. Mol Med 26, 117.

Hengel, H., Hannan, S. B., Dyack, S., MacKay, S. B., Schatz, U., Fleger, M., Kurringer, A., Balousha, G., Ghanim, Z., Alkuraya, F. S. et al. (2021). Bi-allelic loss-of-function variants in BCAS3 cause a syndromic neurodevelopmental disorder. Am J Hum Genet 108, 1069–1082.

Kanca, O., Andrews, J. C., Lee, P. T., Patel, C., Braddock, S. R., Slavotinek, A. M., Cohen, J. S., Gubbels, C. S., Aldinger, K. A., Williams, J. et al. (2019). De Novo Variants in WDR37 Are Associated with Epilepsy, Colobomas, Dysmorphism, Developmental Delay, Intellectual Disability, and Cerebellar Hypoplasia. Am J Hum Genet 105, 413–424.

Knox, E. G., Aburto, M. R., Clarke, G., Cryan, J. F. and O’Driscoll, C. M. (2022). The blood-brain barrier in aging and neurodegeneration. Mol Psychiatry 27, 2659–2673.

La-Vu, M., Tobias, B. C., Schuette, P. J. and Adhikari, A. (2020). To Approach or Avoid: An Introductory Overview of the Study of Anxiety Using Rodent Assays. Front Behav Neurosci 14, 145.

Link, N. and Bellen, H. J. (2020). Using Drosophila to drive the diagnosis and understand the mechanisms of rare human diseases. Development 147.

Liu, N., Schoch, K., Luo, X., Pena, L. D. M., Bhavana, V. H., Kukolich, M. K., Stringer, S., Powis, Z., Radtke, K., Mroske, C. et al. (2018). Functional variants in TBX2 are associated with a syndromic cardiovascular and skeletal developmental disorder. Hum Mol Genet 27, 2454–2465.

Luo, X., Schoch, K., Jangam, S. V., Bhavana, V. H., Graves, H. K., Kansagra, S., Jasien, J. M., Stong, N., Keren, B., Mignot, C. et al. (2021). Rare deleterious de novo missense variants in Rnf2/Ring2 are associated with a neurodevelopmental disorder with unique clinical features. Hum Mol Genet 30, 1283–1292.

Marcogliese, P. C., Shashi, V., Spillmann, R. C., Stong, N., Rosenfeld, J. A., Koenig, M. K., Martínez-Agosto, J. A., Herzog, M., Chen, A. H., Dickson, P. I. et al. (2018). IRF2BPL Is Associated with Neurological Phenotypes. Am J Hum Genet 103, 245–260.

Mohammad, F., Aryal, S., Ho, J., Stewart, James C., Norman, Nurul A., Tan, Teng L., Eisaka, A. and Claridge-Chang, A. (2016). Ancient Anxiety Pathways Influence *Drosophila* Defense Behaviors. Current Biology 26, 981–986.

Moore, B. D., Martin, J., de Mena, L., Sanchez, J., Cruz, P. E., Ceballos-Diaz, C., Ladd, T. B., Ran, Y., Levites, Y., Kukar, T. L. et al. (2018). Short Abeta peptides attenuate Abeta42 toxicity in vivo. J Exp Med 215, 283–301.

Ni, J. Q., Markstein, M., Binari, R., Pfeiffer, B., Liu, L. P., Villalta, C., Booker, M., Perkins, L. and Perrimon, N. (2008). Vector and parameters for targeted transgenic RNA interference in Drosophila melanogaster. Nat Methods 5, 49–51.

Rodriguez, A., Zhang, H., Klaminder, J., Brodin, T., Andersson, P. L. and Andersson, M. (2018). ToxTrac: A fast and robust software for tracking organisms. Methods in Ecology and Evolution 9, 460–464.

Vissers, L., Kalvakuri, S., de Boer, E., Geuer, S., Oud, M., van Outersterp, I., Kwint, M., Witmond, M., Kersten, S., Polla, D. L. et al. (2020). De Novo Variants in CNOT1, a Central Component of the CCR4-NOT Complex Involved in Gene Expression and RNA and Protein Stability, Cause Neurodevelopmental Delay. Am J Hum Genet 107, 164–172.

